# Somatosensory corticospinal tract axons sprout within the cervical cord following a dorsal root/dorsal column spinal injury in the rat

**DOI:** 10.1101/818682

**Authors:** Margaret M. McCann, Karen M. Fisher, Jamie Ahloy-Dallaire, Corinna Darian-Smith

## Abstract

The corticospinal tract (CST) is the major descending pathway controlling voluntary hand function in primates, and though less dominant, it mediates voluntary paw movements in rats. As with primates, the CST in rats originates from multiple (albeit fewer) cortical sites, and functionally different motor and somatosensory subcomponents terminate in different regions of the spinal gray matter. We recently reported in monkeys that following a combined cervical dorsal root/dorsal column lesion (DRL/DCL), both motor and S1 CSTs sprout well beyond their normal terminal range. The S1 CST sprouting response is particularly dramatic, indicating an important, if poorly understood, somatosensory role in the recovery process. As rats are used extensively to model spinal cord injury (SCI), we asked if the S1 CST response is conserved in rodents. Rats were divided into sham controls, and two groups surviving post-lesion for ~6 and 10 weeks. A DRL/DCL was made to partially deafferent one paw. Behavioral testing showed a post-lesion deficit and recovery over several weeks. Three weeks prior to ending the experiment, S1 cortex was mapped electrophysiologically, for tracer injection placement to determine S1 CST termination patterns within the cord. Synaptogenesis was also assessed for labeled S1 CST terminals within the dorsal horn. Our findings show that the affected S1 CST sprouts well beyond its normal range in response to a DRL/DCL, much as it does in macaque monkeys. This, along with evidence for increased synaptogenesis post-lesion, indicates that CST terminal sprouting following a central sensory lesion, is a robust and conserved response.

## 1. Introduction

Spinal cord injury (SCI) affects more than a million people in the United States alone (Reeve Foundation), and has enormous personal and societal impact. Among tetraplegics, the recovery of arm/hand function is cited as the most desired recovery goal for independence and life quality (Anderson, 2004; Anderson et al., 2009), so understanding the neural bases for beneficial changes in hand function is crucial.

The corticospinal tract (CST) is the major descending pathway controlling voluntary hand function in primates (Lemon, 2008; Darian-Smith & Fisher, 2019), and though it plays a lesser role in forelimb / paw function in rodents, it is still important (Anderson et al., 2005; Weishaupt et al., 2013). The CST is conserved across species, but major species differences exist (see Lemon & Griffiths, 2005; Darian-Smith & Fisher, 2019), and in rats brainstem pathways (e.g. the reticulospinal and rubrospinal tracts) are dominant to the CST (Alstermark & Petterson, 2014; Kushler et al., 2002; Morris et al., 2015). As rats are used for the vast majority of SCI studies (Sharif-Alhoseini et al., 2017; Zhang et al., 2014), determining differences/similarities in the recovery process between species is key to assessing the predictive translational power of the rodent model.

In rats the CST originates from two motor (the rostral (RFA) and caudal (CFA) forelimb areas), and two somatosensory cortical regions (S1 and the secondary somatosensory cortex (SII) (Nudo & Masterton 1990; Lemon & Griffiths, 2005). In contrast, in macaque monkeys, it originates from at least 9 different cortical regions (Catsman-Berrevoets & Kuypers, 1976; Dum & Strick, 1991; Galea & Darian-Smith, 1994; Lemon, 2008; Murray & Coulter, 1981), which reflects a more complex brain and sophisticated hand function. Rats are surprisingly dexterous (Steward & Willenburg, 2017), despite having limited fractionation of four usable digits and no digit opposition. However, while both rats and primates can reach and grasp, rat paw performance forms but a small part of the functional repertoire of the highly evolved macaque and human hand.

Though M1 CST sprouting (i.e. from spared fibers) following SCI (Carmel et al., 2013; Jiang et al., 2016; Lindau et al., 2014; Weidner et al., 2001), has long been used as a biomarker of recovery (Kaas et al., 2008; Tuszynski & Steward, 2012), the M1 CST forms only ~ 30% of the total CST fibers in macaques (Darian-Smith et al., 2014; Galea & Darian-Smith, 1994, 1997). Recent work (Darian-Smith et al., 2014; Fisher et al., 2018) in monkeys indicates that the primary somatosensory (S1) CST, which accounts for another ~30% of CST fibers, also plays an important role in the recovery process. In mammalian evolution, the earliest cortical control of movement may well have been from the S1 CST, which suggests this pathway plays an ancient and important role (Lemon & Griffiths, 2005). Following cervical deafferentation injuries that involve the dorsal column, or the dorsal column and dorsal roots (DCL or DRL/DCL), part of the hand is deafferented, and both the motor and somatosensory CSTs sprout (Darian-Smith et al., 2014; Fisher et al., 2018), over the ensuing weeks and months. The S1 CST sprouting is particularly dramatic following a DRL/DCL, and extends bilaterally well beyond the dorsal horn into the intermediate and ventral horns, and caudally as far as T5, or 5 segments (~2.5cm) beyond normal range.

Here we asked if the S1 CST response seen in monkeys is conserved in rats. To address this, we used a combined DRL/DCL model (paralleling recent studies in monkeys) at sub-chronic and chronic time points. The lesion targeted primary afferents innervating the forepaw, and the partially deafferented S1 CST was traced to the cervical cord. A reach-grasp-retrieval task was used to document accompanying recovery in paw function post-lesion. We show in rats that the S1 CST responds to a C6-C7 DRL/DCL injury by sprouting well beyond its normal terminal range, indicating a robust cross species recovery mechanism. We also show accompanying evidence for functional recovery and increased synaptogenesis in S1 CST terminal boutons in the dorsal horn post-lesion.

## 2. Materials and Methods

### Animal model

The rats used in this study were female Sprague Dawleys (250-350g). Animals were individually housed, and on a 12 hour reversed light cycle. They were divided into 3 groups (n=4 per group), including (1) sham controls, (2) short-term sub-chronic animals (rats that survived for 6-7 weeks following the initial lesion), and (3) chronic animals (rats that survived for 10-11 weeks post-lesion). Six-seven weeks post-lesion is considered a transition from the subchronic to chronic phases of recovery when an astrocytic scar has already formed but some immune markers remain in flux (Bowes & Yip, 2014; Kjell & Olson, 2016). Ten-eleven post-lesion weeks is considered a chronic time point in a rat, when functional recovery appears to have plateaued (Bowes & Yip, 2014; Keller et al., 2017; Kjell & Olson, 2016). All lesioned animals received a combined DRL/DCL lesion, since this was previously shown to induce the greatest CST sprouting in monkeys (Darian-Smith et al., 2014; Fisher et al., 2018). Table 1 provides individual animal details. All procedures were performed in accordance with NIH guidelines and approved by the Stanford University Institutional Animal Care and Use Committee.

**Table 1.** Details of the rats used in the current study, showing post-operative survival times, and which rats were used for the confocal immunolabeling analysis. All rats were injected into S1 (D1-3 representation) with the anterograde tracer BDA. *this rat was only used for the confocal imaging analysis.

| ID | Laminectomy to craniotomy (wks) | Post-laminectomy survival time (wks) | Immunolabeling (confocal analysis) |
| --- | --- | --- | --- |
| **Sham-Control** |  |  |  |
| R05 | 4 | 7 | Y |
| R08 | 3 | 6 | Y |
| R14 | 3 | 6 |  |
| R20 | 4 | 7 |  |
| **Post-lesion (6-7 weeks)** |  |  |  |
| R10 | 4 | 7 | Y |
| R15 | 3 | 6 |  |
| R17* | 4 | 7 | Y |
| R29 | 3 | 6 | Y |
| **Post-lesion (10-11 weeks)** |  |  |  |
| R04 | 8 | 11 |  |
| R06 | 8.5 | 11.5 | Y |
| R18 | 7.5 | 10.5 |  |
| R28 | 8.5 | 11.5 | Y |

### Behavior

Rats were tested behaviorally for their ability to perform a reach-grasp-retrieval task, adapted from Schwab (Zemmar et al., 2015), and Whishaw (Whishaw et al., 2008), and modified to better align with the sensory nature of the lesion, and earlier nonhuman primate studies (Darian-Smith & Ciferri, 2005). Rats were acclimated to both handling and to the plexiglass behavior box (15 mins each task per rat/day; 14cm wide × 34cm long × 29cm high; dimensions as per Zemmar et al., 2015; Figure 2).

Rats were placed in the box (Zemmar et al., 2015) for testing, and training sessions were conducted at the same time each day (during the night light cycle). All sessions lasted for 15 minutes or until 20 reach-grasps were recorded (irrespective of success/failure), whichever came first. If 15 minutes passed before 20 grasps were recorded, rats were returned to their home cage. Animals that remained averse to training were removed from further behavioral testing but were handled and fed daily and still used in the study. Cooperative animals were tested and assessed for a minimum of 3 times per week. The task involved pellet retrieval from forceps to best simulate the task used by monkeys. Rats were food restricted as described elsewhere (Zemmar et al., 2015), and weighed daily. Two days prior to surgery, they were given food ad libitum, and post-operatively, rats were closely monitored and again given access to food ad libitum until testing resumed 4-10 days post-surgery (Fornari et al., 2012). Sessions were recorded on a video camera (Canon HD VixiaHF100 Camcorder; 30fps) and analyzed offline using Edius 8 Pro video editing software. The retrieval time (from the time the paw left the window until its return), and success rate of the retrievals were recorded (Figure 2). A reach was considered a success if the animal extended its paw, retrieved the pellet, and was able to eat the pellet, and a fail if the pellet was contacted but not brought to the mouth. Failure to complete the task was interpreted as a deficit in sensory feedback from the digits/paw. Visual feedback was restricted to the reach component of the task, since animals were unable to position their head to see their paw during contact and retrieval. Data were binned per week and plotted (Figure 2), to assess differences in task completion time and success rates, in each paw and pre- and post-lesion.

### Surgical Procedures

All rats underwent a laminectomy, and a craniotomy as described below. Animals were initially anesthetized with 2% isoflurane, and then transferred to a stereotaxic frame and maintained with 1-3% gaseous isoflurane/O_2_ throughout surgery. They were given buprenorphine (0.01-0.05 mg/kg I.P.), atropine sulphate (0.02-0.05mg/kg I.M.), cefazolin (15mg/kg, I.M.), and dexamethasone (1mg/kg, I.M. craniotomy only) prior to surgery. Topical lidocaine (2%), was applied to the ears before placement in the stereotaxic frame, and a line block of bupivacaine (0.25% I.D.) given immediately post-induction. The rat’s respiration and depth of anesthesia were measured every 15 minutes and temperature was continually monitored and controlled using a rodent rectal probe (Braintree Scientific) and thermostatically controlled water heating pad.

At the end of the procedure, the underlying fascia (laminectomy only), and skin were sutured and surgical glue applied externally to seal the skin. Animals were monitored and weighed daily post-operatively, and buprenorphine administered as needed (0.01-0.05 mg/kg I.M.).

#### Laminectomy

Using the C8 spinous process as a landmark, segments C6-8 were exposed dorsally, and for animals receiving a lesion, iridectomy scissors were used to cut dorsal roots C6 and C7 unilaterally (Figure 1). The accompanying dorsal column lesion was made at the C5/C6 border, using a sterile micro scalpel (Graham-Field 2979#15, 15° Angle). The tip of the knife was marked at 1mm to help depth guidance and the incision made into the cuneate fasciculus of the dorsal column. Care was taken to avoid involving the underlying CST. Sham animals underwent the laminectomy, but their dura was not opened. Lesions were made on the side of the preferred paw, wherever a preference could be detected, to help motivate post-lesion use and recovery. Determining a paw preference involved scoring which paw was used the most (>10 times) in 20 reaches (over 2 sessions) to a pellet held in a neutral midline position.

**Figure 1.**
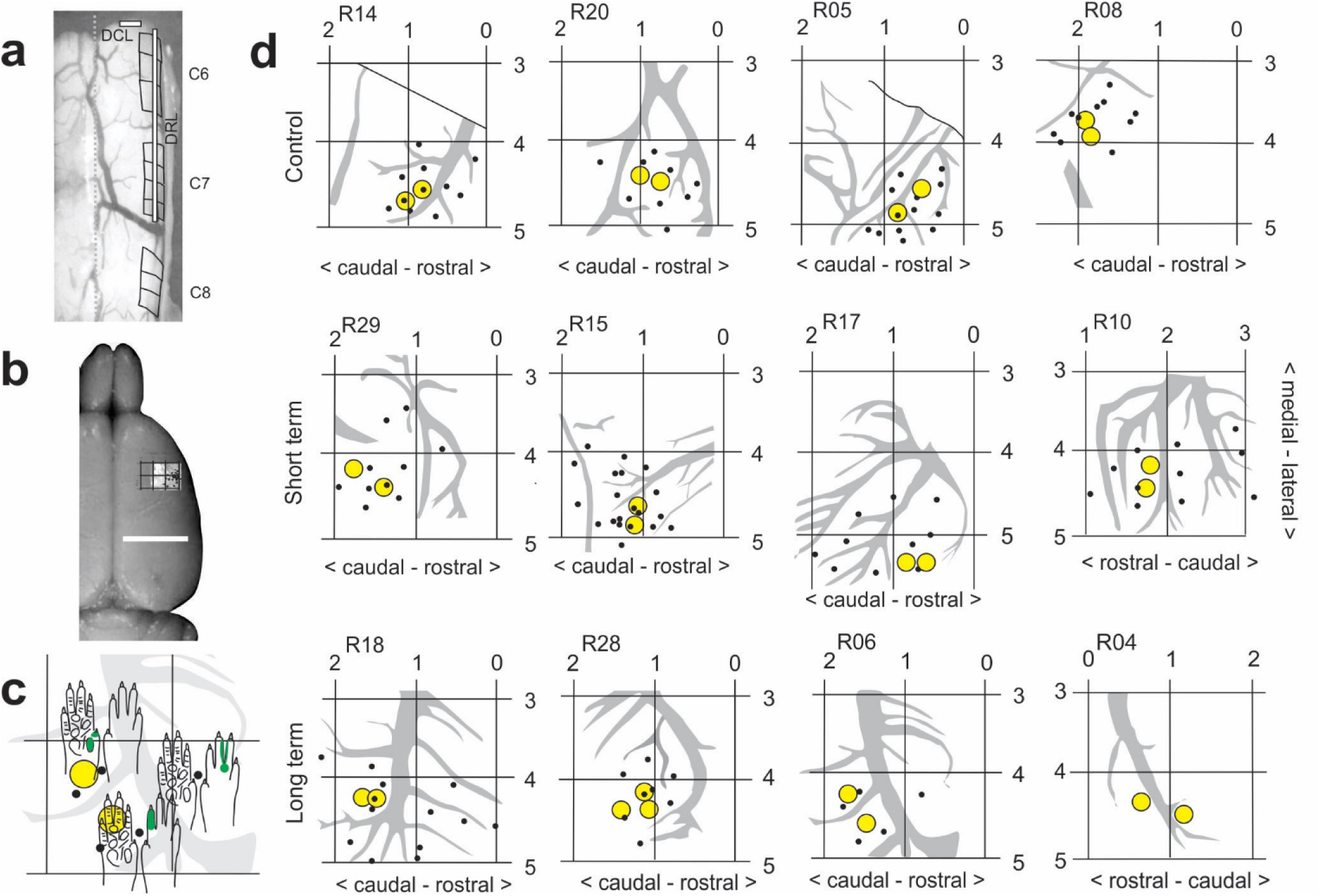
**a**- Dorsal view of the cervical spinal cord during laminectomy, showing an example of the dorsal root and dorsal column lesion placement, which was the same for all lesioned animals. **b**- Dorsal image of rat brain showing location of unitary recordings and forepaw representation in S1. See Neafsey et al., 1986 for classic coordinates of M1 and S1 in rat. **c-d**- Coordinate maps, recording sites (black dots) and anterograde tracer injection locations (yellow circles), made in region of D1-3 representations in S1 in sham-control and lesioned rats. Note that there is greater variability between lesioned animals, presumably due to post-lesion topographic reorganization. **c** shows close up of a part of a cutaneous receptive field (RF) map R06). RFs were mapped for all recording sites (black dots), for tracer placement.

**Figure 2.**
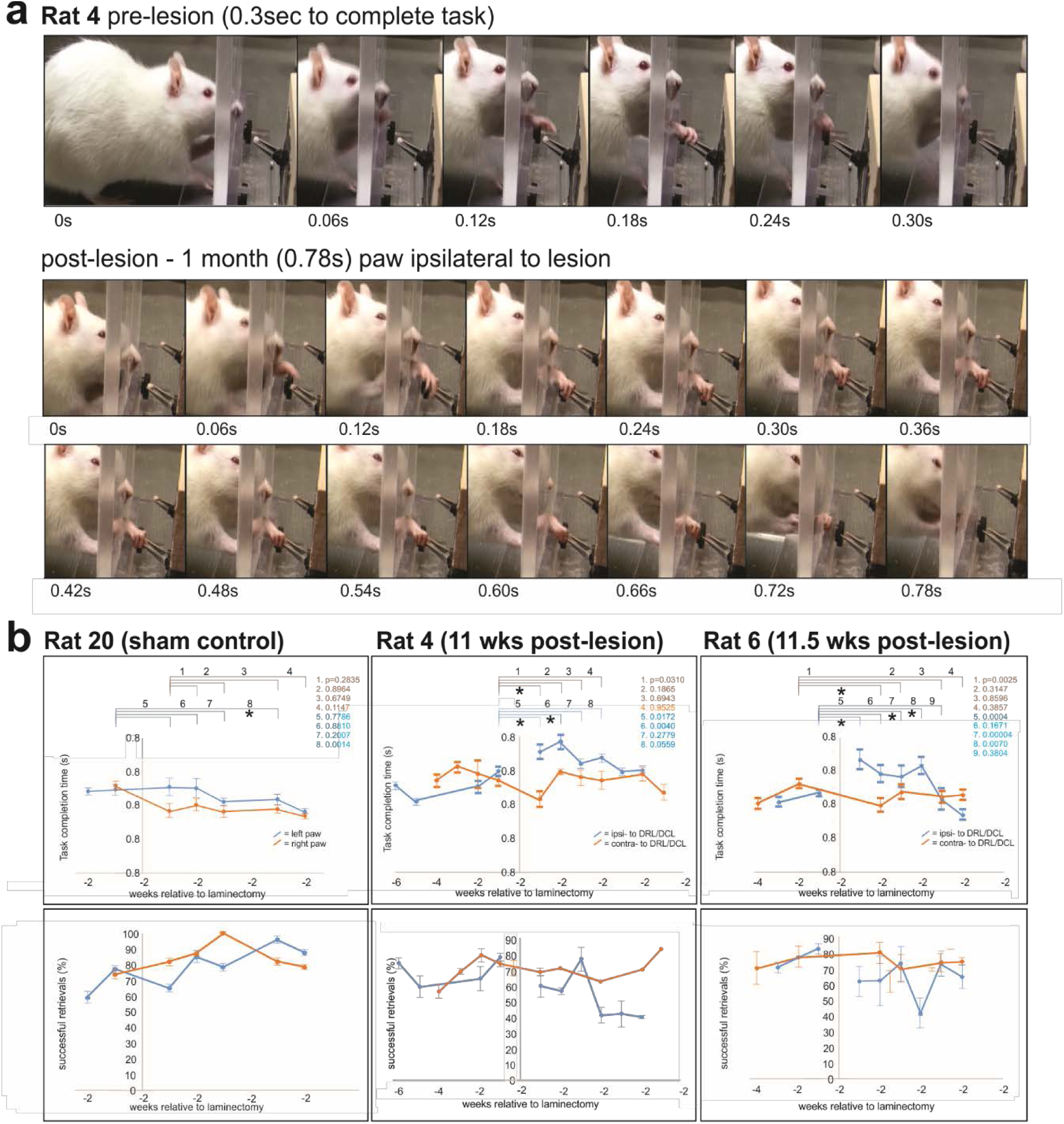
(a) Typical examples of reach-grasp-retrieval sequences obtained before and after a DRL/DCL. (b) shows task completion times (top row), and %age of successful trial (bottom row) data, relative to weeks before and following the laminectomy. Lesioned rats showed an initial increase in task completion times (see Rats 4 and 6), and subsequent improvement over 2-4 weeks, relative to the contralateral paw. This was not observed in sham control rats (e.g. Rat 20), where completion times remained stable for both paws post-laminectomy. In contrast, the proportion of successful trials/week showed no clear trend ipsilaterally following the lesion. (bars = SEMs)

#### Craniotomy

Each animal underwent a unilateral craniotomy, contralateral to the side of the lesion. A small bone flap (approximately 5×5mm) was removed to expose the region of forepaw representation within S1 (relative to bregma: 3-5 mm lateral; 1-3 mm rostral; Figure 1). The dura was resected using iridectomy scissors and electrophysiological recordings made to identify digit representation for anterograde tracer injections.

#### Cortical Recordings and anterograde tracer injections

Electrophysiological (low threshold unitary) recordings were made to locate the region of digit/paw representation within S1. Recordings were made with a tungsten microelectrode (1.2-1.4 mΩ at 1kHz) and peripheral stimulation performed with a camel hair brush and Von Fry filaments as described in earlier studies (Darian-Smith & Brown, 2000; Darian-Smith et al., 2013, 2014; Fisher et al., 2018). The anterograde neuronal tracer biotin dextran amine (BDA, 15% aq., Sigma B9139) was injected into the area corresponding to digit 1-3 representation (D1-3; Figure 1). Injections (0.3ul, 0.8mm deep to pial surface; 2-3 minutes), were made using a glass micropipette (tip <20µm) attached to a Hamilton syringe (constant rate). The overlying skin was then closed.

### Perfusions

Animals were sedated with ketamine (60-80mg/kg, I.P) and euthanized with Beuthanasia (60-80 mg/kg I.P.). Death was confirmed via lack of reflexes (peripheral and corneal), and cessation of breathing. Animals were then transcardially perfused with heparinized 0.1M phosphate buffer (PB) (pH 7.4), and 4% paraformaldehyde in 0.1M PB (pH 7.4, 250ml). The brain and spinal cord were removed and tissue was post-fixed in 4% paraformaldehyde/0.1M PB for 12-24 hours. The brain and spinal cord were carefully dissected, photographed (with roots in place in the spinal cord), blocked and cryoprotected (20% sucrose/dH_2_0). All blocks were snap frozen using isopentane and stored at −80C.

### Tissue processing & analysis

The rostral brain was cut and processed to visualize injection sites, at 50µm, while cord tissue blocks were cut at 40µm. Tissue was cut using a freezing microtome and sections collected into 0.1M PB (pH 7.4). To visualize BDA, sections were reacted with peroxidase (ABC kit; Vector, PK-6100) and the chromagen 3,3’-diaminobenzidine (DAB, Sigma D0426). Tissue was blocked for endogenous peroxidases (Bloxall Vector Laboratories SP-6000), washed in 0.1M PB, incubated in avidin/biotin solution (ABC; Vector PK-6100, 1 h), washed in PB, and incubated with cobalt chloride enhanced DAB, and hydrogen peroxide (0.05%; SigmaFast DAB with Metal Enhancer tablets D0426; 1 tablet each/10mL), in PB. Following a final wash in phosphate-buffered saline (PBS, 0.9% NaCl in 0.1M PB, pH 7.4), tissue was dehydrated and gel mounted (0.5% gelatin in dH_2_O and media from EMS; DEPEX Mounting Media #13514). The injection sites were verified, in sequential cortical sections (50µm at 200 µm intervals).

In the spinal cord, terminal boutons were mapped in coronal section series (every 240µm) through C1-T3, using Neurolucida (MicroBrightField, Inc.). An outline was drawn around the terminal distribution area as described previously (Darian-Smith et al., 2013; Darian-Smith et al., 2014). Results were verified by two separate users blind to the animal test group. Figures were generated using COREL Draw X8.

Immunofluorescence and confocal imaging (Nikon A1R), were used to assess presumed synaptogenesis. A fluorescent tag was used to visualize BDA terminals, and fluorescent secondaries were used to immunolabel synaptophysin (SYN). SYN is a glycoprotein expressed in presynaptic vesicles (Rune et al., 2005) and is used as a marker of synaptogenesis (Masliah et al., 1991). NeuN/MAP2 were combined to label neuronal cell bodies. This allowed us to identify terminals synapsing directly with neuronal soma (see Figure 6), though numbers were too sparse for quantification, and no colabeled BDA/SYN terminals were observed to synapse on to NeuN/MAP2 cell bodies.

For immunofluorescence, sections were washed in phosphate buffered saline with 0.3% triton-X (PBS-TX; 3×10min) and blocked with a 10% solution of normal goat serum (NGS) in PBS-TX (2 hours, room temperature). Primary antibodies (mouse monoclonal anti-NeuN, Millipore-Sigma MAB377, 1:100; mouse monoclonal anti-MAP2, Biolegend 801801, 1:250; and rabbit monoclonal anti-synaptophysin, Invitrogen MA5-14532, 1:500), were diluted in PBS-TX with 3% NGS and sections incubated (48-60 hours at 4 ºC), washed, and exposed to secondary antibodies in PBS (30 min, room temperature; 488 Alexafluor goat anti-rabbit, 1:400, Jackson ImmunoResearch #111-545-003; 647 goat anti-mouse 1:400, Invitrogen A-21235). If BDA was required, sections were washed (PBS) and incubated in ExtrAvidin CY3 (E4142, Sigma Aldrich, 1:200) in PBS overnight. Final washes were with PBS and sections were mounted using 0.5% gelatin and allowed to dry for 45 minutes before coverslipping using antifade mounting media (Prolong Diamond Antifade Mountant, Invitrogen P36965).

### Experimental Design and Statistical Analysis

#### Behavioral analysis

To assess behavioral deficits and recovery in rats, we used two tailed student t-tests (per individual rats comparing week by week as indicated in Figure 2), to determine differences between task completion times before and after laminectomy. Data were obtained for both paws with data from the paw contralateral to the lesion used as an internal control.

#### Terminal bouton distribution analysis

Sections (40 µm) were taken every 240 µm from C1-T3 and the % terminal distribution area/spinal grey area was obtained for each section. The position of each section along the spinal cord was coded according to its vertebral segment, to account for differences in cord length (i.e. total ranged from of 38 to 74 sections per animal). We analyzed data using SAS Studio 3.7 (SAS version 9.04). We coded treatment (Group) as lesion present vs. absent, with type of lesion (short or chronic) nested within. We treated position along the spinal cord as quadratic due to clear non-linearities. Models included all interactions of position and treatment terms. Rat was a random variable nested within treatment. We used repeated measures general linear models (PROC MIXED) to examine % terminal distribution area/spinal grey area.

In keeping with the principle of reduction in animal research, sample sizes were relatively small, yet statistical power was adequate. We tested within-animal effects (position along the spinal cord) using a powerful repeated measures approach that controls for individual differences in average values between rats. Regarding between-animal comparisons (treatment groups), previous research on S1 CST sprouting in nonhuman primates revealed very large effect sizes for the same measures when comparing monkeys between lesion groups (Darian-Smith et al., 2014; Fisher et al., 2018). Our analysis has sufficient power to detect such effects.

#### BDA/SYN colabeled bouton analysis

A similar statistical approach was used to assess changes in numbers of BDA/SYN colabeled S1 CST terminals. Terminal puncta colabeled with BDA and synaptophysin (SYN) were counted within a sample region, selected as the greatest terminal label (i.e. region of greatest reorganization and sprouting) in the medial-ventral region of the dorsal horn. In this analysis, 3 sections were analyzed per segment (C5-C7= 9 sections per animal) in each of 7 rats (see Table 1 and Figures 6–7). As shown in Figure 6, confocal z-stack images were used to identify and count colabeled puncta. In each 40µm thick section, only the middle 20µm region was used in the analysis, to ensure uniformity across sections. Again, we used repeated measures general linear models (PROC MIXED) to examine colabeled bouton counts.

All tests (terminal distribution and synaptogenesis analyses) were two-tailed at alpha = 0.05. We also applied logarithmic or square root transformations in both analyses, as required to satisfy model assumptions (verified by visual inspection of diagnostic plots).

We constructed graphs using the package ggplot2 (Wickham, 2016) for R 3.5.1 (R Core Team 2018).

## 3. Results

A total of 12 female Sprague Dawley rats (250-350g) were used in this study (Table 1). Three weeks prior to experiment completion, a craniotomy was made over the contralateral sensorimotor cortex to expose the S1 cortex. Electrode penetrations/recording sites were marked on photographs of the cortical surface and are shown for individuals in Figure 1. Importantly, stereotaxic coordinates for D1-3 varied by >1.5mm between individual rats of similar size. While this may reflect some topographic reorganization in lesioned animals (Merzenich et al., 1983, 1984), variability was also observed between control animals, where reorganization had not occurred. This indicates normal variations in the population and means that cortical mapping was important for the accurate placement of tracer injections.

Cortical injection sites were posthumously visualized in each rat, in a series of coronal sections (50um) cut through the sensorimotor cortex and processed for BDA, to ensure that tracer injections did not involve subcortical white matter. Since the motor cortex in rats lies only 2-3 mm medial to the S1 paw representation, large injections involving fibers of passage in the white matter could have compromised the data. Such animals were excluded from the study. Dorsal column lesions targeted just the cuneate fasciculus of the sensory dorsal column, and were always made at the C5-C6 border. Though they were not systematically reconstructed in individual rats (due to their location relative to tissue blocking), no change in CST terminal labeling was observed below the level of the DCL. This meant that the CST was not significantly involved by this component of the lesion. The exact mediolateral extent of the DCL was also not critical, since primary afferents traversing this tract were already transected by the DRL, and the central inflammatory reaction was most important to the CST response (Darian-Smith et al., 2014; Fisher et al., 2018).

### Reach-grasp-retrieval task

Though all lesioned rats showed an initial deficit in paw function post-lesion and subsequent recovery in the early post-lesion weeks, the most complete data sets were obtained and analyzed for Rats 20 (Sham), 4 and 6 (chronic lesion animals), and shown in Figure 2. An average of 46 trials were obtained per paw per week in these rats. As shown for lesioned Rats 4 and 6 (Figure 2), task completion times (ipsilateral to the lesion) increased significantly during the early weeks following the DRL/DCL, compared with times in the week prior to the lesion. Times then improved and returned to pre-lesion levels by 3-5 post-lesion weeks. There were no significant changes over time in task completion times for the contralateral paw in sham controls (Rat 20 in Figure 6), where times remained consistent in both paws. Lesioned animals that did not produce full data sets, showed a similar deficit and trend of recovery in the affected paw post-lesion.

In contrast to trial time completion data, no consistent changes were observed in the % age of successful trials per week following the DRL/DCL (see Figure 2). Importantly, rats frequently compensated for the sensory deficit by adapting the placement of their digits and paw during the early post-lesion weeks. This typically involved the transition from approaching the pellet using a lateral – medial sweep, to an approach from above the pellet. In addition, rats used a more pronounced pronation of the wrist in the affected paw post-lesion, and frequently used an over-reach that enabled scooping of the pellet. These behaviors were not observed pre-lesion or contralateral to the lesion.

### S1 CST terminal distributions pre- and post-SCI

In many respects, our results mimic those seen in macaque monkey studies (Darian-Smith et al., 2014; Fisher et al., 2018). In sham control rats, terminal labeling from the S1 CST was confined to a small region of the dorsal horn in the cervical grey matter (Figure 2), in a manner entirely consistent with earlier published work (Flynn et al., 2011). However, following a combined DRL/DCL, input from the S1 CST greatly increased in both the short and long-term lesion groups (Figure 3), indicating that there was significant axon terminal sprouting that extended well beyond normal range. The area of greatest input was located in cervical segments C5-C7 (Figure 4B), or the region of greatest sensory deafferentation following the lesion. There was also a significant rostral expansion of the terminal territory (visible in Figure 3), which coincided with the location of the C3-4 propriospinal network (Flynn et al., 2011; Filli & Schwab, 2015). Figure 4 shows examples of terminal labeling in a sham control and DRL/DCL rats.

**Figure 3.**
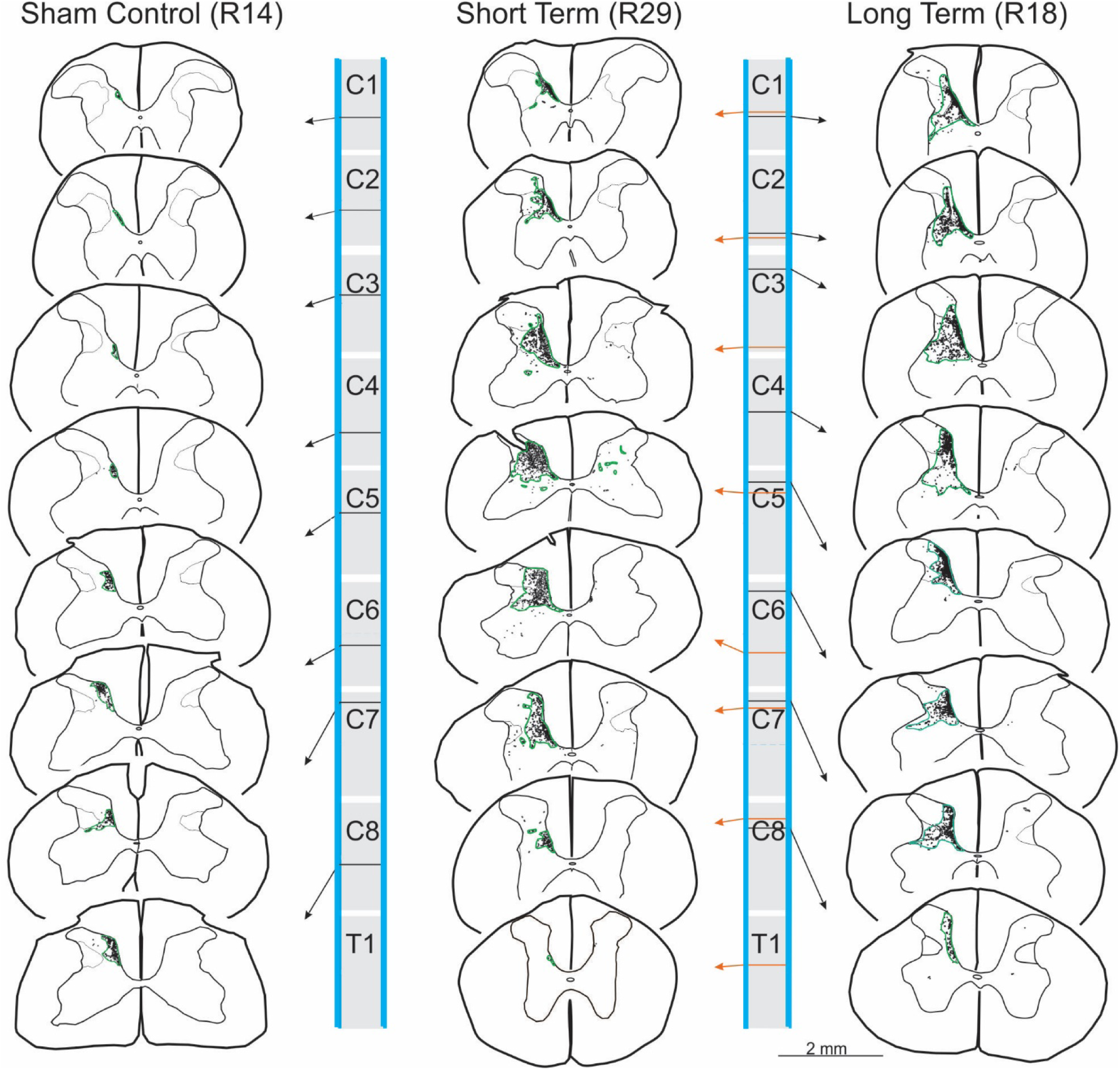
Maps of terminal bouton labeling in spinal cord sections taken from a sample rat from each Group. Green contours show areas of terminal labeling as defined in text. Note the significant expansion of the terminal distribution territory between the sham control and two lesioned groups.

**Figure 4.**
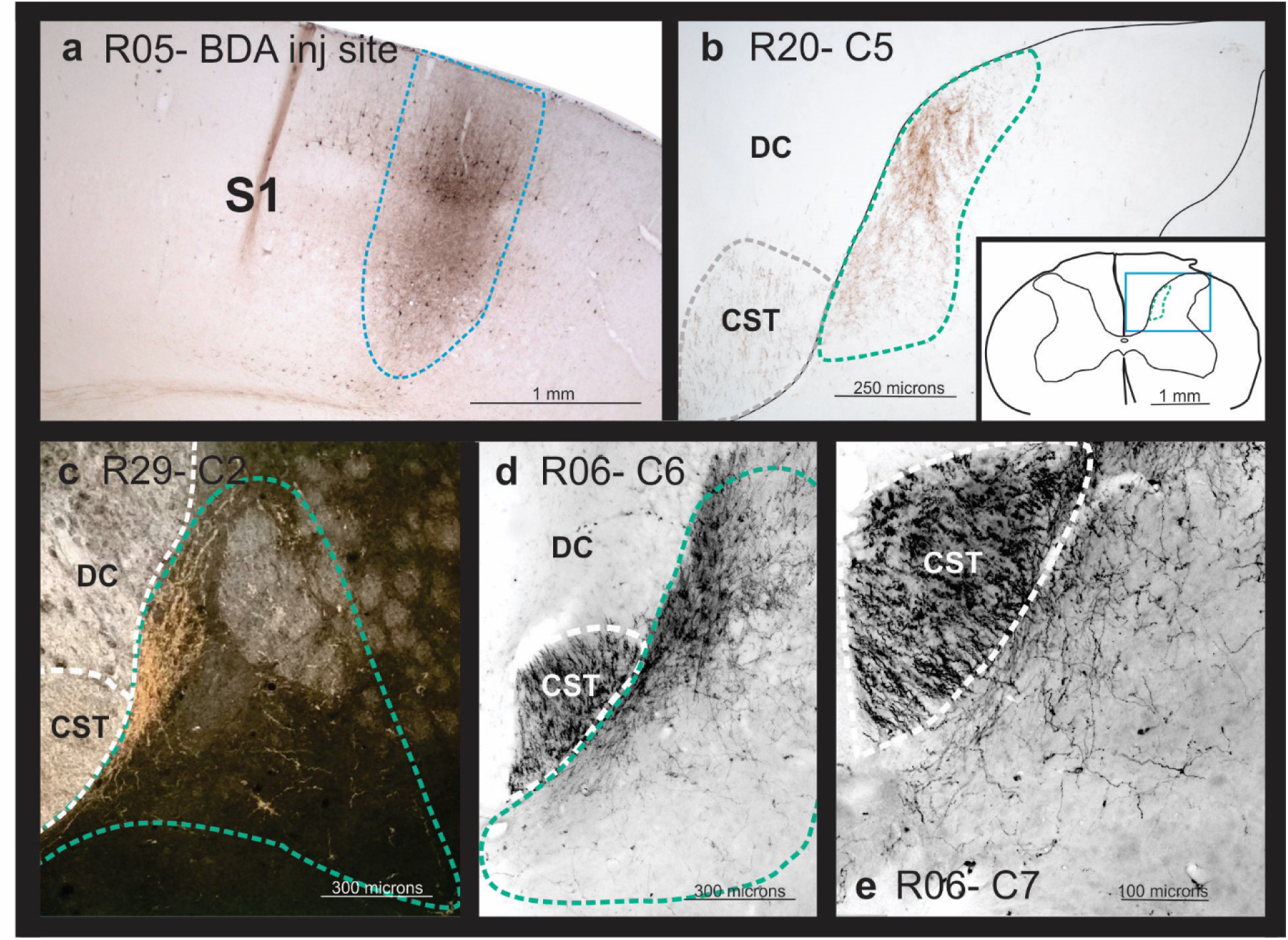
Photomicrographs showing examples of (**a**) a BDA injection site within S1 cortex (within the electrophysiologically identified region of D1-3 representation (shown in Figure 1). **b-e** shows terminal labeling within the dorsal horn in a sham control (**b**) and lesioned (**c-e**) rats. **c** shows a dark field image which highlights the expanded terminal bouton domain and grey matter delineation in this section.

Overall, the % terminal distribution area/spinal grey area on the ipsilateral side showed an inverted-U relationship with position along the spinal cord (Figure 5; quadratic term: F_1,527_ = 203.78, p < 0.0001). Lesioned rats had higher values overall than non-lesioned rats (F_1,8.27_ = 7.79, p = 0.0228), but there was no significant difference between short term and long term animals (F_1,8.24_ = 0.03, p = 0.8616). However, significant interactions between some treatment and position terms (linear and quadratic) show that these differences were not constant across all vertebral segments. The inverted-U relationship peaked at different locations along the spinal cord in lesioned and non-lesioned animals, in lower C4 and lower C5 segments respectively (each segment is one unit long: 4.77 +/− 0.11 vs. 5.65 =/− 0.12, *t* = 5.47, df = 526, p < 0.0001). There was no peak difference between short term and long term lesioned animals (4.88 +/− 0.08 vs. 4.62 +/− 0.18, *t* = 1.33, df = 526, p = 0.185).

**Figure 5.**
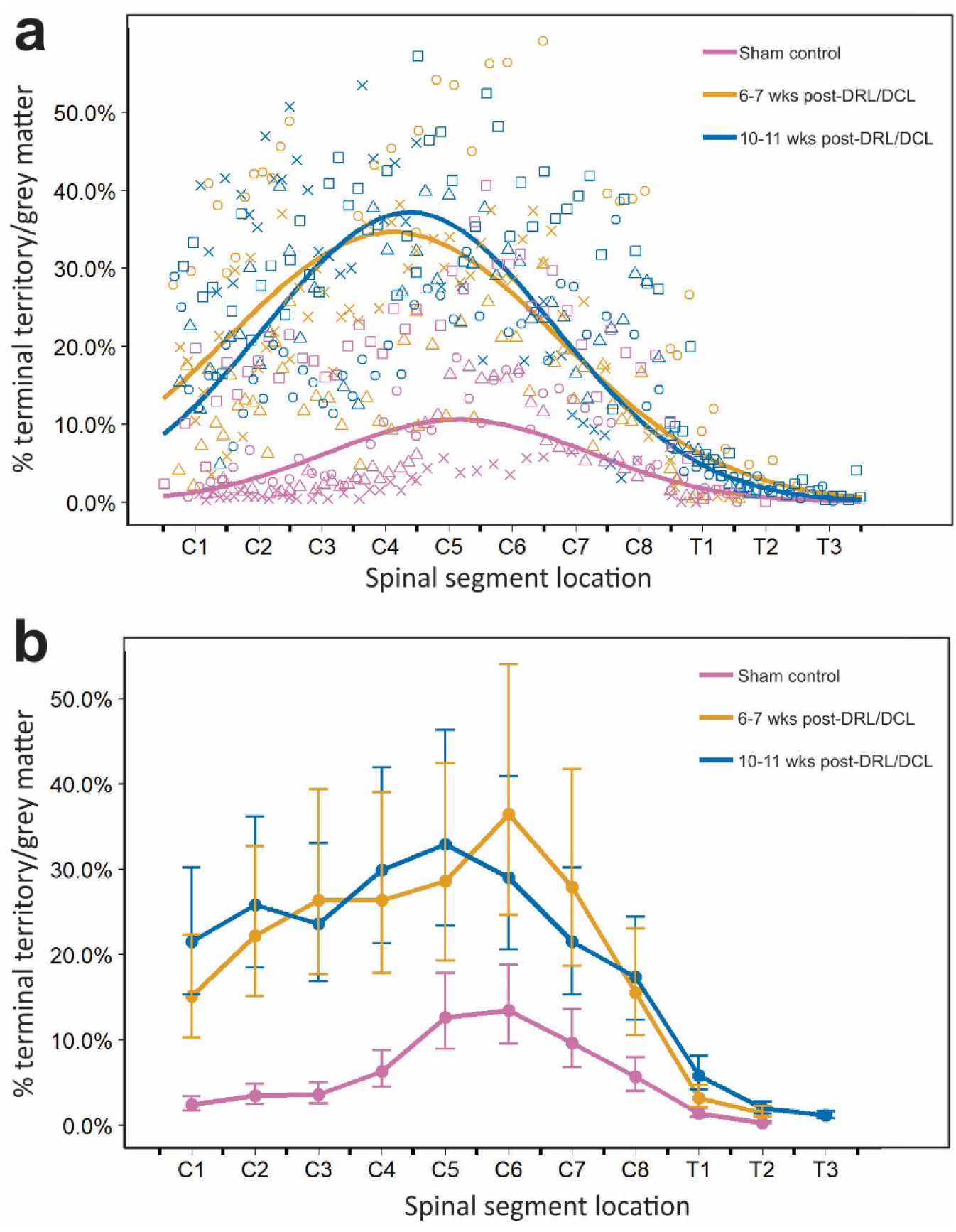
**a,b** Graphs showing percentage of S1 CST terminal bouton territory / total grey matter territory. Both graphs present the same data in two different formats. **a** illustrates raw data for **b**, showing percentage of S1 CST terminal bouton territory (measured by labeled area) / total grey matter territory, versus spinal cord position, for all three groups. The line of best fit was calculated on log-transformed data values (n=2: control (red), short-term (orange), long-term (blue)). The control and lesioned groups are clearly separated, with lesioned animals showing significant terminal sprouting at 6 and 10 weeks post-lesion. The two lesion groups were not significantly different from each other. **b** Graph showing percentage of S1 CST terminal bouton territory / total grey matter territory (least squares means +/− SE; measured by labeled area) versus spinal cord position among all three groups. For ease of visualization, data are binned according to group and vertebral segment and back-transformed for comparison with raw data shown in **a**.

While there is clearly some variability in each of the groups, overall, these data show a significant sprouting of terminal axons in the two lesioned groups compared with the sham control animals. Variability was expected to be greater in rats than was observed in monkeys in our earlier studies (Darian-Smith et al., 2014). This was because we were not able to record from dorsal rootlets prior to making the initial lesion in rats, so lesions were not as specific and individualized as they were in macaque monkeys.

Data were obtained for the tiny amount of terminal labeling observed on the side contralateral to the lesion. However, these data are not reported in detail, as the amount was so small and there were no significant effects of any position along the cord or treatment (lesion and sham groups).

### Evidence of synaptogenesis post-lesion

To determine whether there was evidence for increased synaptogenesis in the S1 CST terminals within the dorsal horn, we examined the relative density of boutons colabeled with BDA and synaptophysin (BDA/SYN), in control versus lesioned rats, in the region of greatest CST terminal labeling in C5-7, in the ventral medial dorsal horn (see Figure 6, and 7).

**Figure 6.**
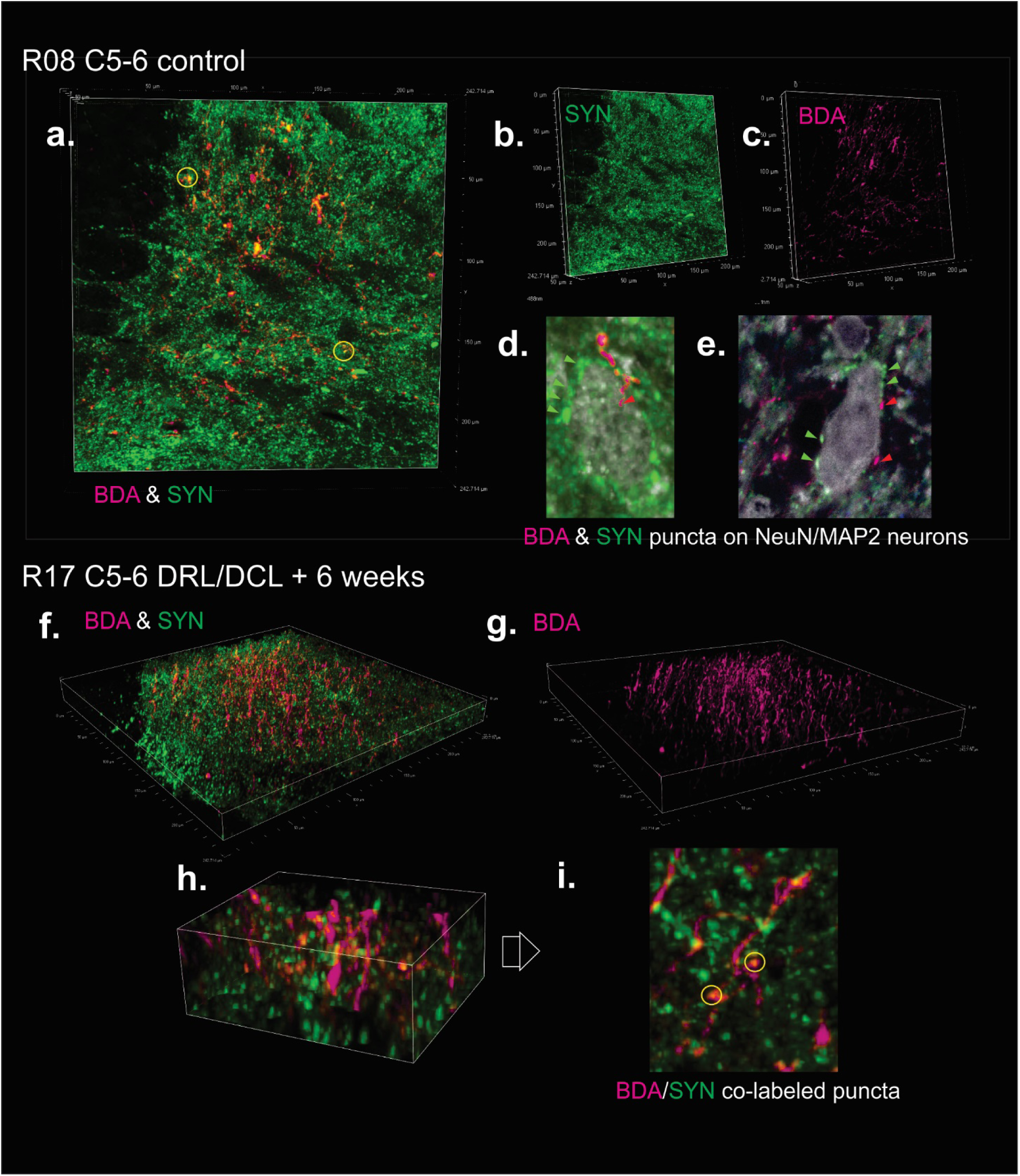
Confocal images in a sham control (**a-e**) and DRL/DCL rat (**f-i**), showing examples of BDA labeled S1 corticospinal fibers and terminals (D1-2 region), and synaptophysin (SYN) labeled terminal boutons. **a-c** and **f-g** show region used for analysis of BDA/SYN colabeling analysis in these sections (see Figure 7). **d-e** show examples of BDA and SYN terminal puncta on to NeuN/MAP2 colabeled neuronal cell bodies. These were observed in all rats, but were sparse, indicating that most synapses were on to dendritic processes. **h**-**i** show localized region of section in **f**, rotated, and with yellow circles indicating BDA/SYN colabeled puncta (i.e. synaptogenesis).

**Figure 7.**
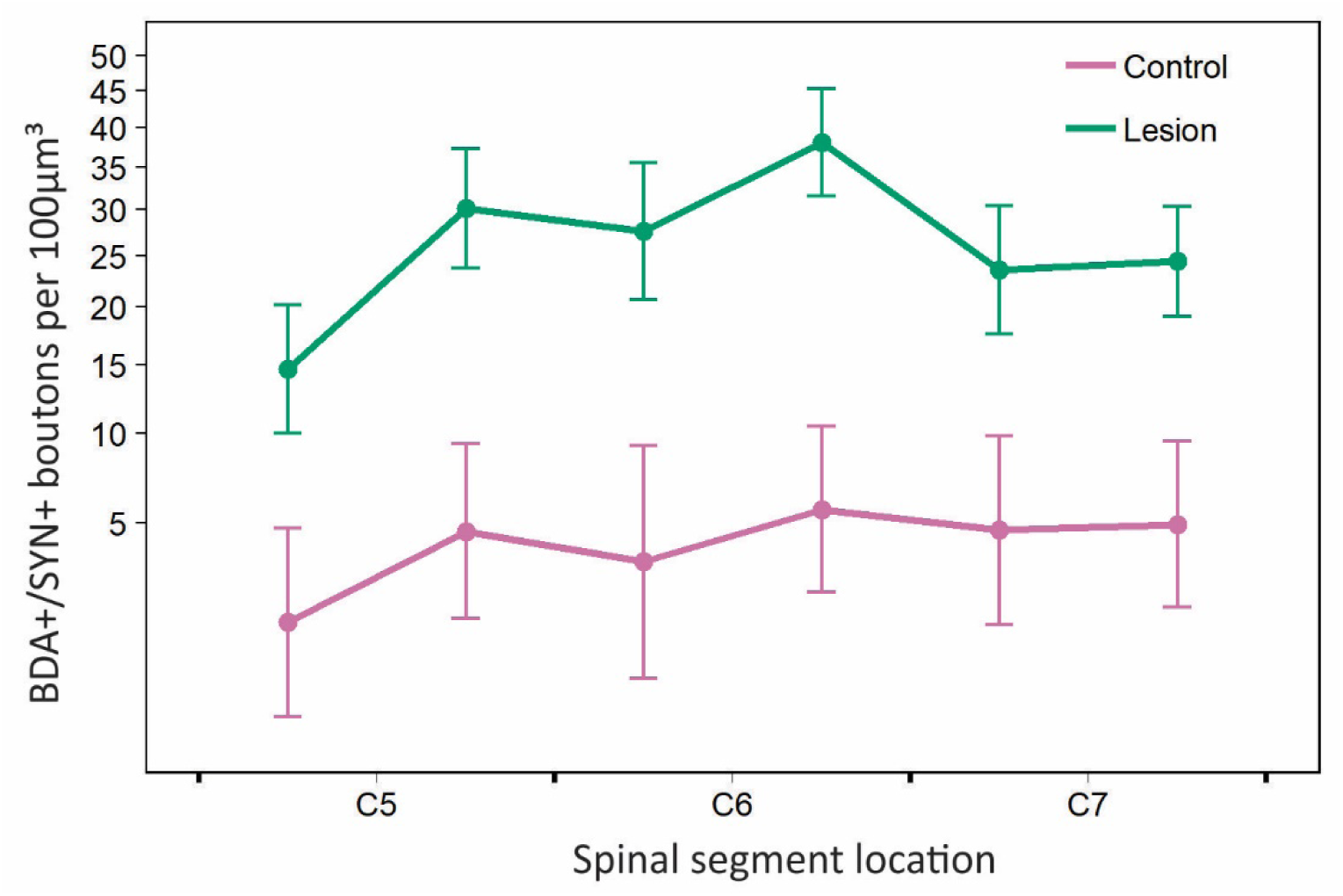
Graph showing BDA/SYN colabeled bouton density (least squares means +/− SE) versus spinal cord position, analyzed from 7 rats (control n=2, short term lesion n=3, long term lesion n=2). Since the short and long term lesion groups had no significant effect on this outcome, the two lesioned groups (see Table 1) were regrouped together. For ease of visualization, data are binned by rostral and caudal halves of each vertebral segment, and the Y axis is transformed accordingly. Least squares means and SEs were obtained from sqrt-transformed data, as in the corresponding statistical analysis.

Colabeled bouton counts showed an inverted-U relationship with position along the C5-7 segments of the spinal cord (Figure 7; quadratic term: F_1,40.1_ = 8.40, p = 0.0060). Lesioned rats had significantly higher counts of BDA/SYN colabeled terminal boutons than non-lesioned (F_1,4.97_ = 13.33, p = 0.0149), but there was no significant difference between short and long term lesion groups (F_1,4.95_ = 1.05, p = 0.3531). There were also no significant interactions between segment position and lesion group variables.

## 4. Discussion

We have shown in the rat for the first time that following a cervical deafferentation (DRL/DCL) spinal injury, the S1 CST sprouts dramatically within the spinal cord (C1-T1), well beyond its normal territory. This is of interest because it means that the S1 CST response is robust across mammalian species. The implication is that this is an important response mechanism that contributes to the recovery of paw and hand function (in skilled reach-grasp behaviors), following spinal injuries that involve primary afferents. Our findings in the rat model suggest that terminal sprouting is needed for circuitry rewiring to occur, but they do not indicate a clear correlation between the extent of CST terminal sprouting and the extent of behavioral recovery. The details of this relationship are far more nuanced and will require targeted future research to clearly ascertain (see Darian-Smith et al., 2014). Understanding this mechanism and relationship is, however, of fundamental importance for developing strategies that optimize neuronal reorganization that best promotes functional recovery.

Our findings show that there was a significant expansion in the S1 CST terminal bouton distribution in rats receiving a combined DRL/DCL compared with sham control animals. No differences, however, were found between the 6-7 and 10-11 week post-lesion periods, which means that the terminal growth had largely occurred within the preceding weeks. If consolidation of these projections over time does occur and if it involves pruning of the exuberance to something approaching pre-lesion ranges (Lorenzana et al., 2015; Humanes-Valera et al., 2017; Nakanishi et al., 2019), this was not evident at 11 post-lesion weeks.

An initial deficit was observed in the rat’s ability to reach, grasp and retrieve a food pellet, such that rats took significantly longer to complete the task post-lesion, and showed a general clumsiness. Detectable behavioral recovery occurred within the first 2-5 weeks, and compensatory strategies were adopted, including a greater paw pronation, and tendency towards using a dorsal overreach for scooping the target. Similar deafferentation injury models in nonhuman primates (Darian-Smith & Ciferri, 2005; Darian-Smith et al., 2013, 2014; Fisher et al., 2018; Crowley et al., unpublished) similarly show a deficit and recovery of digit use over weeks and months post-lesion.

Presumed synaptogenesis (Masliah et al., 1991), also increased during this early post lesion period to significantly above baseline, which supports the idea that many of the newly formed terminal sprouts form functional synapses with postsynaptic targets.

The general pattern of terminal sprouting observed in both control and lesioned rats is similar to that observed previously in macaque monkeys (Darian-Smith et al., 2014). Similarities even extend to exuberant sprouting in the rostral cervical cord (see Figure 3). This is where cervical (C3-C4) propriospinal networks (Flynn et al., 2011; Taccola et al., 2017 for review), have been implicated in SCI recovery and hand dexterity (Tohyama et al., 2017; Isa, 2019), though the relationship between these neurons and the S1 CST terminal sprouting has yet to be determined.

### Variability among animals

There was considerable variability of terminal distribution patterns between animals in this study. This is likely due to normal inter-animal variation, as well as to the limitations of the rat lesion model. Given that rats are small, it was not feasible to perform electrophysiological recordings in dorsal rootlets, prior to making the lesion (Darian-Smith et al., 2014). As a consequence, C6 and C7 rootlets were cut (since they have known input from the forepaw), and the central injury was always made at the C5/6 border. This did not allow for small inter-animal differences and the exact rootlets through which sensory input from specific digits travelled, could not be definitively determined in the rodent model. This means that while all the injections made during the craniotomy were accurately localized to the region of D1-3 representation, these would not have always labeled the epicenter of CST axons most affected by the DRL/DCL.

### Dorsal horn circuitry and the role of S1 CST terminal sprouting in recovery

Following the DRL/DCL used in this study, primary afferent inputs were removed from C6-C7 segments. This permanently disrupted the circuitry in the medial ventral dorsal horn, which makes this region a focal point for synaptic reorganization. The S1 CST terminal growth in the dorsal horn was likely enabled by the inflammatory response caused by the central dorsal column component of the DRL/DCL (Bollaerts et al., 2017). This was evident in monkey studies comparing peripheral and central spinal injuries (Darian-Smith et al., 2013, 2014; Fisher et al., 2018).

In normal rats, as in primates, S1 CST terminals synapse in the (contralateral) dorsal horn primarily on to the presumed axons/dendrites of intraspinal neuronal populations (Schieber, 2007; Gosgnach et al., 2017; Ziskind-Conhaim & Hochman, 2017; Côté et al., 2018). Target neurons may include local reflex, long and short range propriospinal (Filli & Schwab, 2015) and commissural interneuronal populations (Bannatyne et al., 2003; Jankowska et al., 2009; Soteropoulos et al., 2013), which in turn connect with motor neurons, and/or regulate (or gate) ascending tracts (i.e. the dorsal columns, spinocerebellar and spinothalamic pathways). Recent studies in rodents (and particularly in mice), underscore just how complex and multi-functional some of these interneuronal populations are (Del Barrio et al., 2013; Bourane et al., 2015; Abraira et al., 2017; Ueno et al. 2018). However, little to nothing is known of their involvement in adaptive rewiring post-SCI.

The axon growth observed in this study, may be the most efficient way for spared S1 axons to reach and establish new functional circuitry. And, it is easy to speculate that the S1 CST’s role in regulating afferent flow, provides a strong adaptive drive to establish connections that re-stabilize this information flow. Importantly, monkey work shows that following a DRL (Darian-Smith et al., 2004) or similar DRL/DCL (Fisher et al., unpublished), spared primary afferents also sprout within the dorsal horn (and cuneate nucleus where S1 cortical input also projects). Assuming that primary afferents also sprout post-lesion in the rat, we speculate that the axon terminal sprouting response is widespread, and that it occurs throughout the neuraxis wherever there are spared inputs, and wherever normal information flow is disrupted.

### Species differences

Though the S1 CST terminal sprouting observed in this study after a DRL/DCL was broadly similar to observations made in macaque monkeys (Darian-Smith et al. 2014), significant species differences exist and deserve consideration (Darian-Smith & Fisher, 2019; Ebner and Kaas, 2015; Walcher et al., 2018). Terminal sprouting overall was less dramatic in the rat than the monkey. For example, the S1 CST (from digit region) did not extend caudally beyond T1 in the rat post-lesion. In contrast in monkeys, following a similar DRL/DCL, both the M1 and S1 CSTs projected 3-4 segments beyond their normal range. This reflects a species difference, since it cannot easily be explained by differences in the methodology of the two studies.

Rodents, like primates, have more mechanoreceptors in the glabrous skin of their forepaws compared with their hindpaws (Walcher et al., 2018), which reflects the use of the forepaw in tactile exploration (Steward & Willenberg, 2017; Kameda et al., 2019). However, macaque monkeys and higher primates have evolved a vastly more sophisticated hand/digit function. Primate evolutionary success has depended on precision pincer grasp and fine manipulation capabilities, thumb opposition, and independent digit use. In all, hand-brain coevolution in primates (including a vastly expanded prefrontal cortex), has enabled a primate hand dexterity that dwarfs that of the rat forepaw.

In rats, the CST has 4 functional subdivisions, while this number more than doubles in macaques and higher primates in step with their larger brain. In rats, more so than in primates, the bulbospinal projections (e.g. rubrospinal and reticulospinal) play a major role in the regulation of directed forepaw motor control (Brosamle & Schwab, 1997; Lemon, 2008). For example, the rubrospinal tract projects directly onto motoneurons innervating muscles of the distal forelimb, and its ablation in rodents affects the animals’ paw rotation as well as their accuracy in judging target distance during fine motor tasks (Kuchler et al., 2002; Morris et al., 2015). In macaque monkeys, in contrast, the CST is critical to volitional hand/digit function (Galea & Darian-Smith, 1997; Lemon & Griffiths, 2005), and the rubrospinal tract plays a lesser role (Burman et al., 2000; Lemon, 2016; Garcia-Alias et al., 2015). In humans the trend continues, so that the rubrospinal tract is all but inconsequential.

In macaque studies, following a DRL/DCL, the S1 CST input was observed to expand bilaterally (Darian-Smith et al., 2014). In this study, tracers were only injected contralateral to the lesion site, so it was not possible to determine the extent of sprouting from the ipsilateral cortex. However, it seems likely that rats also experience a bilateral expansion of the S1 CST post injury, given our findings and the species similarities on the side of the lesion. Further work will be needed to clarify this phenomenon in the rodent so that its potential role in recovery can be assessed.

### Translational value of the rodent model in spinal injury?

A major motivation of the present study was to determine the suitability of the rat as a translational model for the investigation of corticospinal tract changes following spinal injury. Our findings indicate that despite the many differences between the rodent and primate species, the S1 CST sprouting response mechanism to the spinal injuries modeled in this study, are clearly robust and largely conserved. The implication is that the rat SCI model described here, presents a useful translational tool, for addressing basic questions about post-injury CST sprouting, that cannot readily be addressed in the clinically more relevant nonhuman primate model.

## Acknowledgements

This work was supported by the National Institute of Neurological Disorders and Stroke (R01 NS091031 to CD-S), and the Stanford Department of Comparative Medicine MLAS program. We would like to thank Joe Garner for his early guidance with statistics.

## Data Availability Statement

The data that support the findings of this study are available from the corresponding author upon reasonable request.

